# Millennia of genomic stability within the invasive Para C Lineage of *Salmonella enterica*

**DOI:** 10.1101/105759

**Authors:** Zhemin Zhou, Inge Lundstrøm, Alicia Tran-Dien, Sebastián Duchêne, Nabil-Fareed Alikhan, Martin J. Sergeant, Gemma Langridge, Anna K. Fotakis, Satheesh Nair, Hans K. Stenøien, Stian S. Hamre, Sherwood Casjens, Axel Christophersen, Christopher Quince, Nicholas R. Thomson, François-Xavier Weill, Simon Y.W. Ho, M. Thomas P. Gilbert, Mark Achtman

## Abstract

*Salmonella enterica* serovar Paratyphi C is the causative agent of enteric (paratyphoid) fever. While today a potentially lethal infection of humans that occurs in Africa and Asia, early 20^th^ century observations in Eastern Europe suggest it may once have had a wider-ranging impact on human societies. We recovered a draft Paratyphi C genome from the 800-year-old skeleton of a young woman in Trondheim, Norway, who likely died of enteric fever. Analysis of this genome against a new, significantly expanded database of related modern genomes demonstrated that Paratyphi C is descended from the ancestors of swine pathogens, serovars Choleraesuis and Typhisuis, together forming the Para C Lineage. Our results indicate that Paratyphi C has been a pathogen of humans for at least 1,000 years, and may have evolved after zoonotic transfer from swine during the Neolithic period.

**One Sentence Summary:** The combination of an 800-year-old *Salmonella enterica* Paratyphi C genome with genomes from extant bacteria reshapes our understanding of this pathogen’s origins and evolution.

## Main Text

According to historical records (*1*), humans have long been afflicted by bacterial infections. Yet comparative genomics of extant bacterial pathogens routinely estimate a time to the Most Recent Common Ancestor (tMRCA) of no more than a few centuries (*2*). In contrast, tMRCAs of millennia were estimated by other analyses that included ancient DNA (aDNA) (*3*,*4*). One possibility to account for these apparent discrepancies is that the tMRCA of extant isolates only includes the currently existing crown lineage, and ignores older sub-lineages that are now extinct. To test the generality of this inference, we scanned metagenomic sequences from teeth and long bones of 33 skeletons who were buried in Trondheim, Norway between 1100-1670 CE (*5*) (Fig. 1A, C). 266 *Salmonella enterica* reads were identified in one tooth from skeleton SK152. We reconstructed a genome (designated Ragna) from that *Salmonella* after additional metagenomic sequencing of nine libraries from SK152 derived from one bone, two teeth and dental calculus (table S1). We then provided an evolutionary framework for invasive salmonellosis, and its host specificity, by analyzing the ancient Ragna genome (fig. S1) within the context of ~50,000 modern genomes from *S. enterica* which we had assembled with the <u>EnteroBase</u> online resource (*6*).

**Fig. 1.**
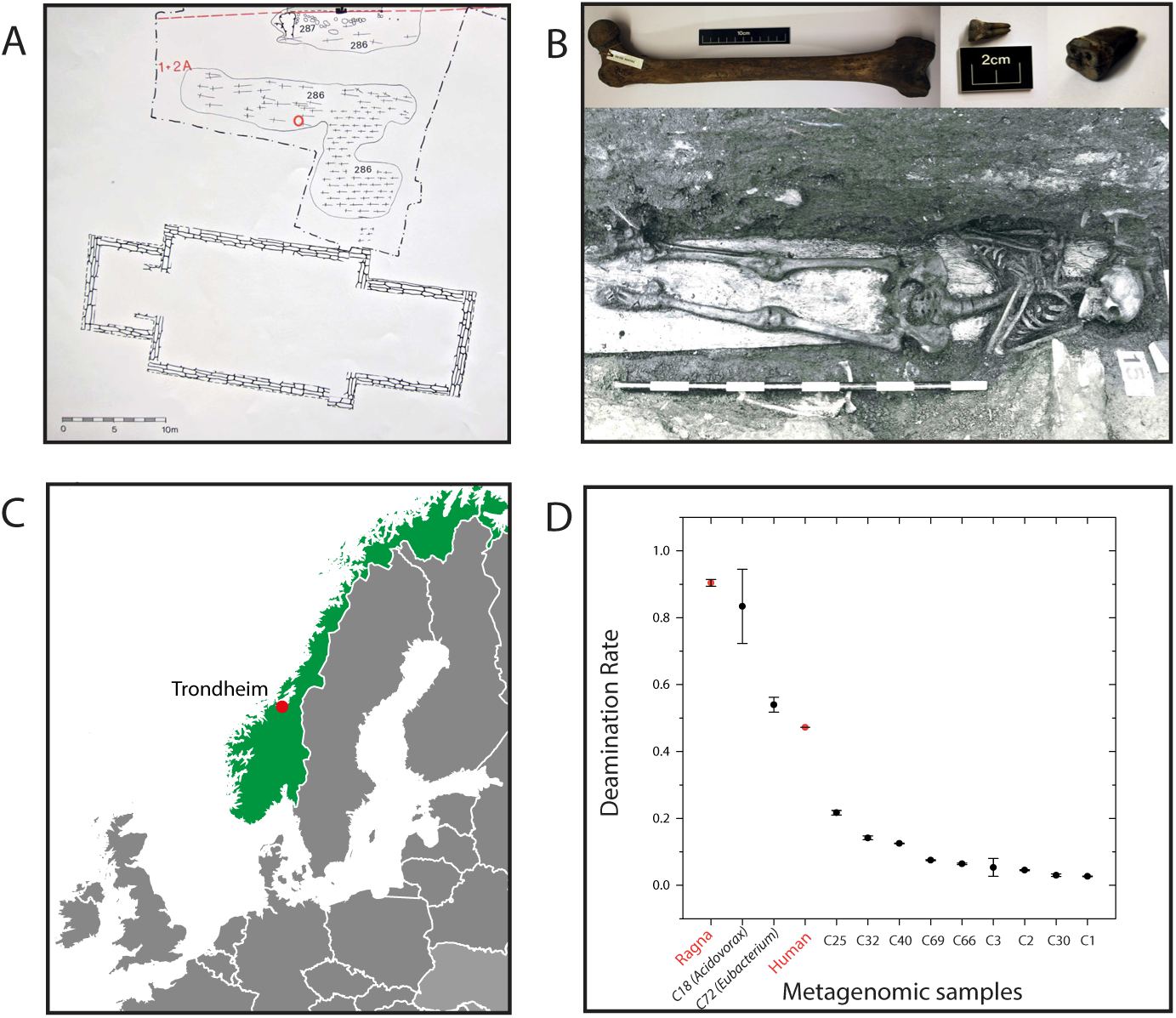
Geographic, archaeological and metagenomic features of skeleton SK152. A) Excavation site (Folkebibilotekstomten, 1973-1985) of the church cemetery of St. Olav in Trondheim, Norway. The burial location of SK152 (red circle) belongs to a building phase which has been dated archaeologically (5) to 1200 CE (range 1150-1250). B) Femoral long bone plus two teeth from which *Salmonella* DNA was extracted (Top) and entire skeleton (bottom). C) Map of Europe surrounding Norway (green). D) Deamination rate for metagenomic reads in the *Salmonella* Paratyphi C Ragna genome, human DNA and 11 sets of assemblies identified by CONCOCT (34) (C1 through C72). The high deamination of C18 (*Acidovorax*) and C72 (*Eubacterium*) shows that they are as old as reads from humans or Ragna while the other assemblies likely represent modern environmental bacteria.

SK152 is the skeleton of a 19-24 year old woman of 154±3 cm height who was buried in 1200±50 CE according to archaeological investigations (*5*). Calibrated radiocarbon (^14^C) dating of two teeth estimated her burial as 100 or 200 years before this (fig. S2), a discrepancy that can be explained by the reservoir effect on radiocarbon dating when fish products were the predominant diet (*7*). Based on δ^18^O_carbon_ isotopic measurements from her first and third molars, this woman likely migrated from the northernmost inland areas of Scandinavia or Northwest Russia during her childhood, and arrived in Trondheim by her early teens (table S2). The skeleton rested on a wooden plank (a symbolic coffin) in a grave filled and covered with anoxic, acidic and waterlogged wood chips and soil (Fig. 1B).

From the metagenomic reads, we also succeeded in reconstituting 11 clusters of contigs that correspond to nearly complete microbial genomes (Fig. 1D; fig. S3; tables S3-S4). Nine of these genomes likely reflect recent (*8*) soil contamination because their 5’-single-stranded deamination rates were low (DNA damage <0.22) *versus* values of 0.47 for human DNA and 0.9 for Ragna (table S5). Two of the nine genomes are from bacterial sulfate-reducers and a third consists of archaeal methanogens, typical of the microbes that would be common in the soil surrounding SK152. In contrast, the two other assembled genomes are likely to have been endogenous to this corpse since burial because they exhibited high levels of DNA damage. C72, a novel species of *Eubacterium*, was found almost exclusively in dental calculus (table S6), and may have been a major component of a biofilm associated with periodontal disease, as are other taxa of *Eubacterium* (*9*). C18 belongs to *Acidovorax*, which is associated with plant pathogens (*10*), and may have been introduced with the wood chips. None of the few assembled contigs related to *S.enterica* were greater than 1 kb in length, and we therefore reconstructed the Ragna genome by read mapping against its close relatives.

In order to identify close relatives of the Ragna genome, we first created phylogenetic trees of 2,964 genomes from EnteroBase that represent the genetic diversity of *S. enterica* subspecies *enterica*. We used the 711,009 SNPs (single-nucleotide polymorphisms) from 3,002 concatenated core genes (2.8 Mb) from these genomes (table S7) to infer a maximum-likelihood tree (Fig. 2A). We also inferred an independent species tree from gene-by-gene alignments of each core gene. The topologies of the two trees were identical near the root and towards the tips, and at 75% of the intermediate branches. These trees were used for phylogenetic placement (MGPLACER) of the metagenomic reads from SK152; all reads specific to *S. enterica* were assigned to a monophyletic clade of serovar Paratyphi C genomes. That clade was in turn contained within the “Para C Lineage”; which includes related monophyletic clades of serovars Choleraesuis, and Typhisuis (*11*), plus one genome of the extremely rare serovar Lomita. We used two genomes of serovar Birkenhead as an outgroup for rooting trees of the Para C Lineage because they were the closest genetic relatives (Fig. 2).

**Fig. 2.**
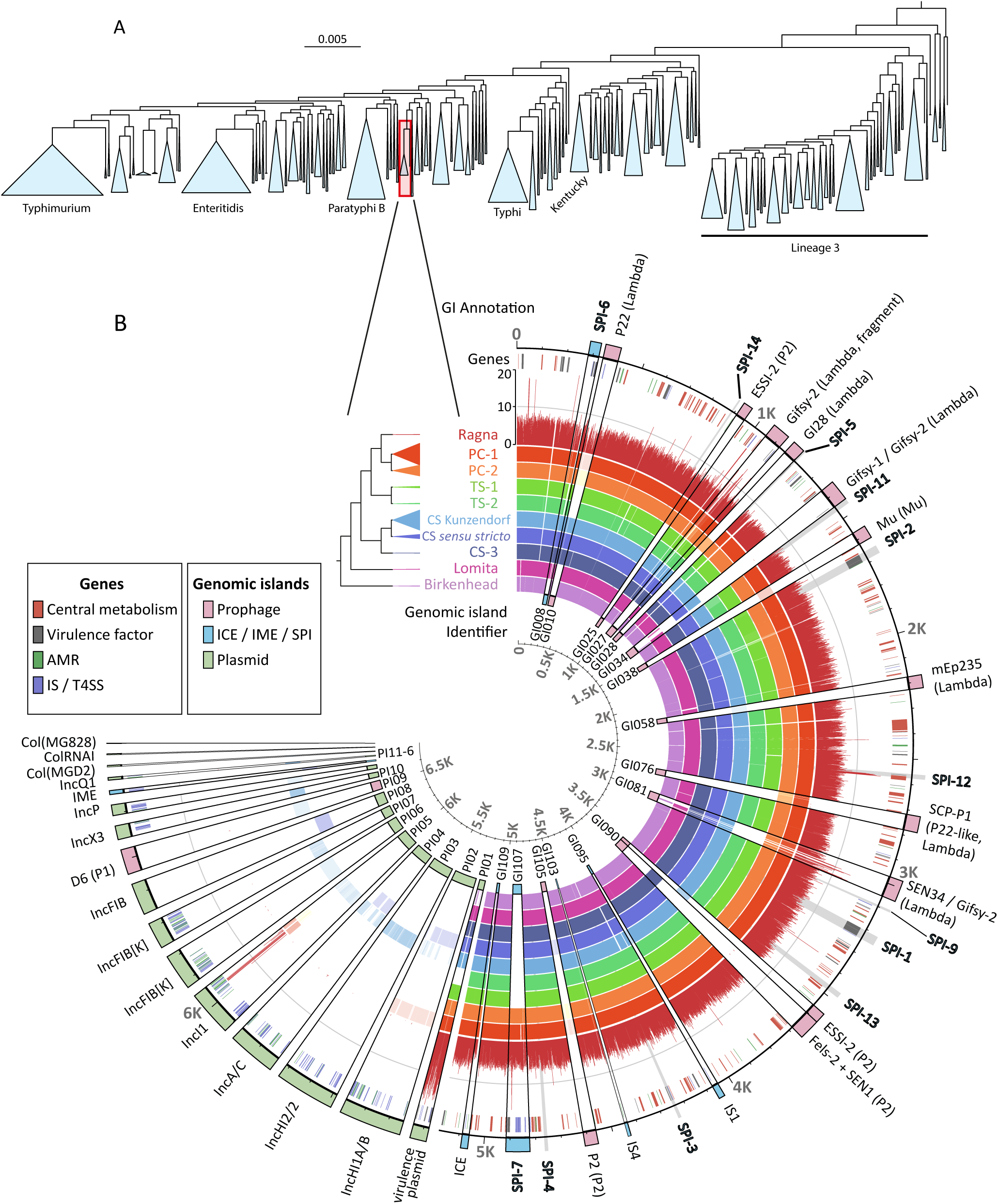
Genomic phylogenies of *S. enterica* and the Para C Lineage. A) Maximum-likelihood phylogeny of representative genomes of *S. enterica* subspecies *enterica*. Each genome is a random representative of one of the 2,964 sequence types that had been defined by ribosomal multi-locus sequence typing (rMLST). The Para C Lineage and other lineages containing numerous genomes are collapsed into blue triangles, and common serovars that are associated with several of those lineages are indicated under the tree. Red rectangle: Para C Lineage plus Birkenhead. B) Pan-genomic contents for 220 genomes of the Para C Lineage, including Ragna. Genes are ordered as in Database S1 (primary ticks every 500 genes). Circles (inner to outer): Circle 1. Sixteen major, variably present chromosomal genomic islands (GI008-GI109; Database S2) followed by sixteen cytoplasmic elements (PI01-PI16), color-coded as in the right key. Circles 2-10: The frequency of presence or absence of each gene per sub-lineage within the phylogram is indicated by color opacity. Circle 11: Coverage of aDNA reads per gene within Ragna (Scale 0-20 at 12:00). Circle 12: Genes color-coded as in the left key. Circle 13: Traditional annotations of GIs, PIs and other invariably present genomic elements, such as SPIs (gray wedges).

Paratyphi C was first recognized in 1916 when it was isolated from Eastern European soldiers suffering from enteric fever (also called paratyphoid fever) (*12*). Enteric fever is a severe, occasionally fatal, illness that was also common at that time in northern Africa, South America and North America. It has since disappeared from Europe, except for occasional travelers returning from East Asia or Africa (*11*). The Ragna genome demonstrates that Paratyphi C bacteria were present in northern Norway 800 years ago, and the presence of Ragna sequences within both teeth and bones in a young adult suggests that SK152 died of septicemia associated with enteric fever. Paratyphi C only infects humans but other members of the Para C Lineage show different host specificities. Serovar Choleraesuis is associated with septicemia in swine (and occasionally humans) and Typhisuis with epidemic, chronic paratyphoid/caseous lymphadenitis in domestic pigs (*11*, *13*). Although both serovars continue to cause disease in southern and eastern Asia, Choleraesuis is rare in Europe today, except in wild boar (*14*), and Typhisuis has been eradicated from European pigs through culling of infected herds.

The inference of tMRCA and phylogeographic history is best performed with bacterial strains that were collected from global sources over a broad temporal range. However, only 108 of >50,000 assembled *Salmonella* genomes in EnteroBase belonged to the Para C Lineage, and their geographical and temporal sources were very limited. We therefore combed the strain collection at the Institut Pasteur, Paris, for historical Para C Lineage strains from diverse sources, and sequenced 111 additional genomes, which resulted in a total of 219 modern Para C Lineage genomes from multiple continents and dated from 1914-2015 (table S8). These genomes defined sub-lineages within each of the serovars (Paratyphi C: PC-1, PC-2; Choleraesuis: CS Kunzendorf, CS *sensu stricto,* CS-3; Typhisuis: TS-1, TS-2) (Fig. 2B). The metagenomic reads from SK152 were then mapped (after initial processing and de-duplication) against all 219 genomes in order to identify all reads mapping to the pan-genome of the Para C Lineage. We found 1,030,108 unique Para C Lineage-specific reads in teeth (0.05-0.18% of all reads) or the femur (0.01%), but not in dental calculus (table S1). The Ragna reads covered 98.4% of a reference Paratyphi C genome (RKS4594) with a mean read depth of 7.3-fold. In order to avoid spurious SNP calls associated with DNA damage, we only called SNPs in the Ragna genome against the 95% of RKS4594 that was covered by at least two reads.

The evolutionary dynamics and selective pressures associated with local ecological interactions are often thought to result in variation of gene-content in microbes (*15*). We therefore anticipated that 800 years of evolution would have resulted in dramatically different gene content between the Ragna and modern Paratyphi C genomes, and expected even greater differences from the other serovars within the Para C Lineage. Surprisingly, this was not the case - of the 4,388±99 genes (total length 4.8±0.08 Mb) in a Para C Lineage genome, 78% were intact core genes (table S9, Fig. 2B), and only 604 core SNPs distinguished Ragna from the MRCA of modern Paratyphi C (Fig. 3). Some of the core genes are universally present in the Para C Lineage plus Birkenhead, even though they belong to mobile genetic elements that are variably present in other *Salmonella*, e.g. the pathogenicity islands SPI-1 to SPI-4, SPI-6, SPI-9 and SPI-11 to SPI-14. Similarly, the virulence plasmid was present throughout the Para C Lineage except for Typhisuis sub-lineage TS-2. A further constant feature of the Para C Lineage was the absence of genes encoding typhoid toxin, which is thought to trigger enteric fever by serovars Typhi and Paratyphi A (*16*).

**Fig. 3.**
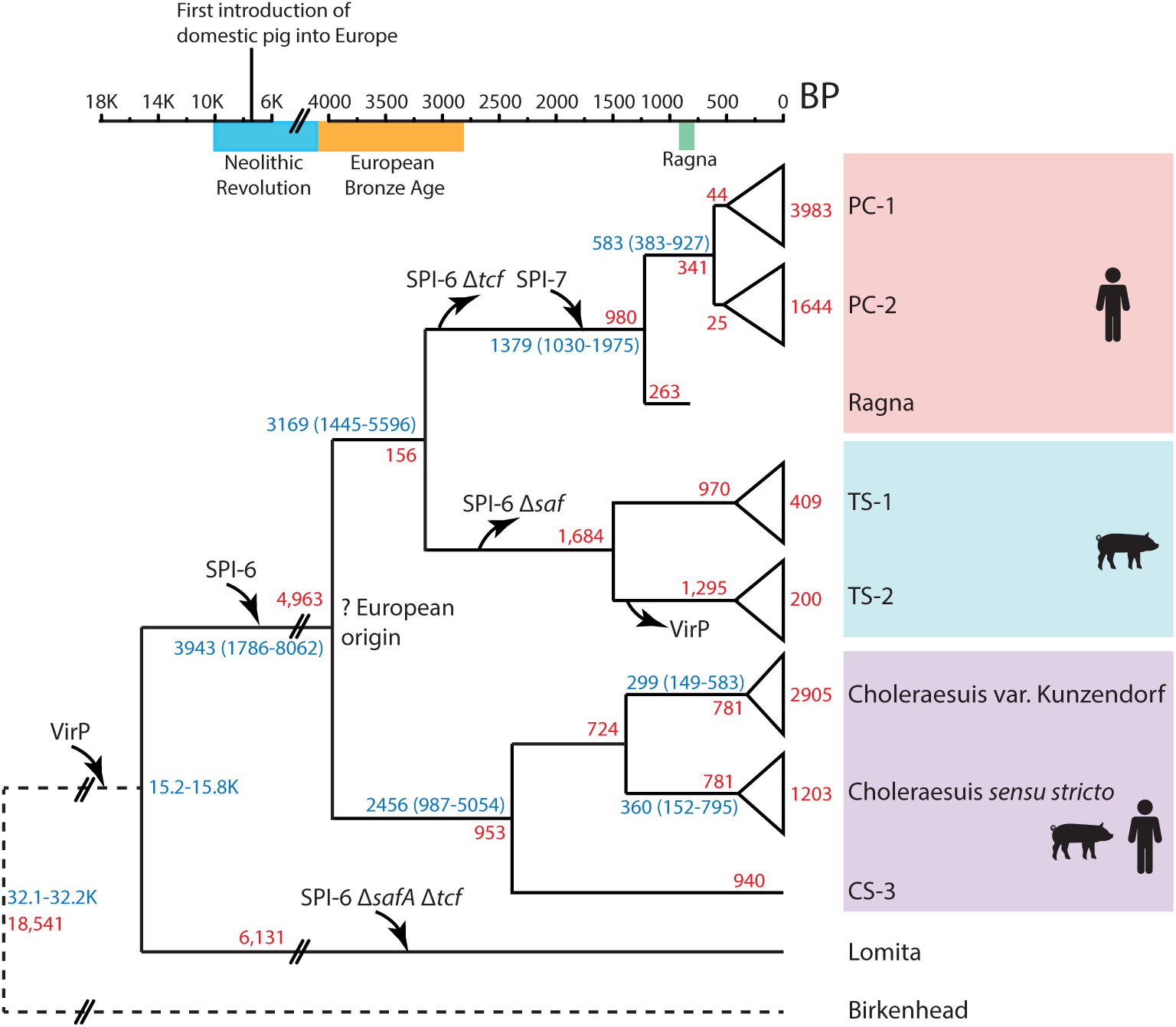
Cartoon of the evolutionary history of the Para C Lineage on the historical timeframe BP for human history in Europe (top). The phylogram indicates the acquisition of the virulence plasmid (VirP), SPI-6 and SPI-7 by inward arrows and deletions of parts of SPI-6 by outwards arrows (see fig. S6). Blue numbers on branches indicate the average of the median date from two independent subsamples plus the 95% extreme credible intervals of both samples (tables S11-S13). Red numbers indicate numbers of non-recombinant core single nucleotide variants. The host specificity of the individual serovars is indicated at the right.

Other studies have indicated that microbial host adaptation is accompanied by the accumulation of pseudogenes (*17*). Pseudogene accumulation is thought to be due to the streamlining of genes that are no longer necessary for the infection of multiple hosts (*18*), but has also been proposed to provide an efficient mechanism for rewiring of transcriptional regulation (*19*). A median number of 40-60 pseudogenes was observed within the 2,964 representative *Salmonella* genomes. Twenty-five pseudogenes were inferred for the MRCA of the Para C Lineage and 69 for the MRCA of Paratyphi C, Choleraesuis plus Typhisuis (fig. S4), suggesting that the MRCA had not yet adapted to a particular host, or had only recently begun to do so. However, the splits into the individual serovars may well mark the beginnings of host adaptation because higher numbers of pseudogenes were inferred for the MRCAs of each of the serovars (Choleraesuis: 95; Paratyphi C: 116; Typhisuis: 181), and the number of pseudogenes continued to increase to modern times (fig. S4). It is possible that some of these pseudogenes contribute to host specificity, rather than simply representing functions that are not required for infection of those particular hosts.

We also attempted to identify mobile genetic elements that could account for the different host specificities of the individual serovars among the 3,901 genes in the accessory genome of the Para C Lineage. These accessory genes clustered together within 227 GIs (mobile elements and genomic islands), including 37 plasmids, 32 prophages, 16 IMEs (Integrative and Mobilizable Elements), SPI-5 to SPI-7, and two ICEs (Integrative and Conjugative Elements) (table S10). The GIs were acquired or lost on 311 independent occasions, of which at least 60% are unlikely to be important for host specificity because they were restricted to a single genome (table S10). Most of the other gains or losses are also unlikely to represent successful evolutionary changes in virulence or host specificity, because, as in other *Salmonella* (*20*–*22*), they were restricted to individual sub-lineages within a serovar, and sister sub-lineages differing in the possession of those genes are also prevalent in invasive disease (Fig.2B, fig. S5, table S11). For example, the integration of a P22-like prophage designated SCP-P1 (GI076), upstream of *pgtE* in Paratyphi C has been reported to prevent opsonization via the PgtE omptin (*23*) and drastically reduce the minimal infectious dose for laboratory infections of mice (*24*). However, GI076 is absent from half of Paratyphi C genomes, including Ragna (Fig. 2B). Similarly, the *fimH102* allele of a type 1 fimbrial adhesin was reported to endow specific adhesion to porcine cells on serovar Choleraesuis (*25*). However, the entire *fimH* gene is absent in CS *sensu stricto*. Indeed, none of the virulence factors and GIs seemed likely to be consistently related to differential virulence or host specificity (table S11), with the notable exceptions of SPI-7 and SPI-6.

SPI-7 (GI107) is a pathogenicity island in serovars Typhi, Paratyphi C and Dublin which encodes the Vi capsular polysaccharide. Vi prevents the opsonization and clearance that is triggered by binding of the C3 component of complement to lipopolysaccharide (*26*), and might thereby promote enteric fever in humans. Our data showed that SPI-7 is present in all genomes of Paratyphi C, including Ragna, but absent from the other serovars in the Para C Lineage, suggesting that SPI-7 was acquired prior to the expansion of serovar Paratyphi C, and might be associated with host specificity. A 5-gene deletion within SPI-7 was present throughout sub-lineage PC-1, but this does not affect the production of the Vi polysaccharide (*11*).

SPI-6 (GI008) is present throughout the entire Para C Lineage. It encodes a Type Six Secretion System (T6SS) as well as *Salmonella* atypical fimbriae (*saf*) and Typhi colonization factor (*tcf*) (figure S6). T6SS systems encode intracellular, inverted bacteriophage-tail-like structures that can inject lethal effectors (TaeX) into proximal eukaryotic and bacterial cells (*27*). For *S.enterica* serovar Typhimurium, SPI-6 contributes to gastrointestinal colonization and pathogenesis in mice (*28*) and chickens (*29*). Expression of *tcf* in *Escherichia coli* resulted in specific adhesion to human epithelial cells (*30*) and *saf* mutations within serovar Choleraesuis reduced gastrointestinal colonization of pigs (*31*). All T6SS genes within SPI-6 were present throughout the Para C Lineage but *tssL*, *tssM* and *vgrG* were absent in the outgroup, serovar Birkenhead, suggesting that the acquisition of an intact T6SS by horizontal gene transfer (HGT) during the evolution of the ancestor of the Para C Lineage may have facilitated its proficiency at gastrointestinal colonization and causing invasive disease. Interestingly, SPI-6 varied between the different serovars within the Para C Lineage (figure S6), and the various SPI-6 variants might also be associated with the differential host specificity of the serovar. Precise independent deletions between pairs of 43 bp-long, identical direct repeats within SPI-6 have cleanly excised the *tcf* genes within serovars Lomita and Paratyphi C, including Ragna, and the *saf* genes in serovar Typhisuis. SPI-6 in Lomita may represent an independent acquisition by HGT: The termini of the deletion of *saf* genes in Lomita differed from the termini in Typhisuis, and a Tae4/Tai4 toxin/antitoxin T6SS effector is found exclusively in Lomita’s SPI-6. Furthermore, unlike other members of the Para C Lineage, the SNP density against a reference genome of Choleraesuis was fourfold higher (0.015) in SPI-6 than in the core genome. Thus, a parsimonious progression of these events would be an initial acquisition of a large SPI-6 by HGT after Lomita branched off, followed by successive deletions prior to the MRCAs of Paratyphi C and Typhisuis. However, alternative evolutionary paths in which each SPI-6 variant represents an independent HGT event cannot be excluded.

These observations immediately raised the question of evolutionary time-scales. How old are the individual serovars? And do these genomes allow an estimate of the minimal age for the infection of warm-blooded animals by *S. enterica*?

We dated key stages in the evolution of the Para C Lineage using a Bayesian phylogenetic approach (Fig. 3, tables S12-S13), and confirmed by randomization of the dates of isolation (figure S7) that there was a strong temporal signal in two individual sub-samples of each dataset. The mean molecular clock rate was the same (9.4×10^-8^ substitutions per site per year; CI: 6.9×10^−8^ - 1.2×10^−7^) for Paratyphi C whether Ragna was included or excluded for these calculations (table S12). However, even though the dates of isolation of the extant bacteria spanned almost 100 years, and they had been obtained from multiple continents, analyses that excluded Ragna estimated the mean tMRCA as 583 y (CI: 383-927) whereas those that included the Ragna genome estimated 1,379 y (CI: 1,030-1,975) (table S13). Using the somewhat different age estimates for the Ragna genome provided by archaeological reconstructions (1200 CE) or C^14^ dating (1073 CE) had little influence on the estimated tMRCA (table S12). Thus, adding a single aDNA genome from an extinct sub-lineage to 219 genomes from a broad sample of extant bacterial isolates increased the tMRCA by twofold.

The maximum age of Paratyphi C is 3,169 y, the date when Paratyphi C split from its closest relatives, serovar Typhisuis. In turn, the estimated tMRCA for Paratyphi C, Typhisuis plus Choleraesuis was 3,943 y, and that ancestor evolved in Europe according to three independent Bayesian and maximum-likelihood methods (figure S8, table S14). We note that a European ancestry is consistent with the existence of Ragna in northern Norway 800 years ago, and of other Paratyphi C genomes in Mexico in 1,545 CE that may have been imported to the New World by Europeans (*32*). We also note that serovars Choleraesuis and Typhisuis infect swine, and that the tMRCA dates overlap with the Neolithic domestication of pigs from wild boar in Europe (*33*) (Fig. 3). It is therefore possible that this invasive sub-lineage evolved in swine or in boar, and that Paratyphi C represents a subsequent host adaptation to humans by a zoonotic pathogen of domesticated animals. This proposal is eminently testable through examining aDNA from boar and swine in Europe for genomic traces of DNA related to the Para C Lineage.

We calculated increasingly more speculative estimates of the dates of earlier branch points (table S13) by extrapolating the inferred mutation rates from the Bayesian analyses to the species tree estimated by maximum likelihood. The estimated tMRCA of the entire Para C Lineage including Lomita was ~15.500 y and the split from Birkenhead was ~32,000 y. However, because these estimates are extrapolations over long time periods, they are best considered to represent ballpark figures until substantiation has been provided by additional aDNA analyses. It is not currently feasible to distinguish mutational SNPs from recombinational SNPs at the species level in organisms which recombine as much as *S. enterica* We therefore simply note that the estimated age of *S. enterica* subspecies *enterica* was ~70,000 y on the basis of all core SNPs, in order to provide an initial, flawed estimate of its age that will need correction.

Our analyses indicate that numerous recent estimates of tMRCAs based on comparative genomics of extant bacteria will need revision as additional genomes are derived from aDNA studies. Our results also indicate that both the core and accessory genome of bacterial pathogens are unexpectedly stable over millennia, and that much of the dramatic variation between extant genomes represents transient genetic fluctuation whose evolutionary relevance to ecological fitness is uncertain. Finally, these analyses provide a paradigm for how combining genomes from both extant and ancient bacteria can facilitate the reconstruction of their evolutionary history.

## Acknowledgments

All genomes referred to here are available from the publicly available workspaces entitled “<u>rST representatives</u>” (2964 genomes) and “<u>Para C Lineage</u>” in the *Salmonella* database in EnteroBase (http://EnteroBase.warwick.ac.uk). Interactive versions of Fig. 2 can be found at https://enterobase.warwick.ac.uk/anvi_public/ParaC_pangenome. Data supplements are permanently stored at the University of Warwick (http://wrap.warwick.ac.uk/85593) and non-human metagenomic data at https://sid.erda.dk/wsgi-bin/ls.py?share_id=E56xgi8CEl. EnteroBase (BBSRC BB/L020319/1) was developed by N-F.A., M.J.S. and Z.Z. (equal contributions) under guidance by M.A. Additional grant support was from the Wellcome Trust (202792/Z/16/Z), Danish National Research Foundation (DNRF 94) and Lundbeckfonden (R52-5062). For other attributions see Supplementary Material.

